# Infectious SARS-CoV-2 is emitted in aerosols

**DOI:** 10.1101/2021.08.10.455702

**Authors:** Seth A. Hawks, Aaron J. Prussin, Sarah C. Kuchinsky, Jin Pan, Linsey C. Marr, Nisha K. Duggal

## Abstract

Respiratory viruses such as SARS-CoV-2 are transmitted in respiratory droplets and aerosols, which are released during talking, breathing, coughing, and sneezing. Non-contact transmission of SARS-CoV-2 has been demonstrated, suggesting transmission in aerosols. Here we demonstrate that golden Syrian hamsters emit infectious SARS-CoV-2 in aerosols, prior to and concurrent with the onset of mild clinical signs of disease. The average emission rate is 25 infectious virions/hour on days 1 and 2 post-inoculation, with average viral RNA levels 200-fold higher than infectious virus in aerosols. Female hamsters have delayed kinetics of viral shedding in aerosols compared to male hamsters, with peak viral emission for females on dpi 2 and for males on dpi 1. The majority of virus is contained within aerosols <8 µm in size. Thus, we provide direct evidence that, in hamsters, SARS-CoV-2 is an airborne virus.

## Introduction

SARS-CoV-2 is a respiratory virus that has caused more than 190 million cases and 4.1 million deaths, as of July 2021 [1]. Respiratory droplets and aerosol particles (aerosols), which may contain virus, are expelled during coughing, sneezing, talking, and breathing and can vary widely in size from less than 1 µm to greater than 100 µm [2, 3]. Their size significantly impacts transmission risk and mode due to differences in the way that droplets and aerosols travel through the air. There is a continuum of maximum distances that particles can reach. Those smaller than 10 μm remain suspended in air for many minutes to hours, during which they can travel long distances; this does not rule out their potential to transmit at close range, too.

Infectious SARS-CoV-2 has been cultured from aerosols sampled near COVID-19 patients [4-7]. SARS-CoV-2 has also been isolated from aerosols <1 µm within a car driven by a COVID-19 patient with mild illness [8]. The collection of exhaled breath condensate (EBC) is a non-invasive sampling method of respiratory droplets and aerosols that has been used to assess the airborne transmission potential of respiratory viruses, including seasonal human coronaviruses, influenza viruses, and rhinoviruses [9-12]. For COVID-19 patients, the emission rate of SARS-CoV-2 RNA in EBC was estimated to be >1 million viral RNA copies/hour by breathing [13, 14]. In non-human primates, SARS-CoV-2 RNA has also been detected in EBC collected from inoculated animals [15, 16].

Hamsters are a naturally-susceptible animal model for SARS-CoV-2 transmission that develop few clinical signs of disease [17-22]. Unlike mice, hamsters model asymptomatic infection, which has been suggested to be the most important component of community transmission of SARS-CoV-2 [23, 24]. Oral swabs from inoculated hamsters contain high levels of infectious virus, with levels similar to saliva collected from COVID-19 patients [25, 26]. Importantly, inoculated hamsters have been shown to transmit SARS-CoV-2 to naïve hamsters via non-contact transmission [17, 18, 22], suggesting SARS-CoV-2 may be transmitted between animals via aerosols released during breathing. However, infectious SARS-CoV-2 has not yet been cultured from aerosols released from infected animals.

In this study, we seek to determine the shedding kinetics of infectious SAS-CoV-2 in aerosols. We find that hamsters inoculated with SARS-CoV-2 emit infectious SARS-CoV-2 into the air prior to and concurrent with the onset of clinical signs of disease. Viral titers decrease quickly over time, whereas viral RNA is detected in the air for many days post-inoculation. Male hamsters shed virus in the air earlier than female hamsters. We also find that aerosols <8 µm contain the majority of airborne virus.

## Materials and Methods

### Viruses and cell lines

Inoculations were performed using SARS-CoV-2 strain USA-WA1/2020 (BEI Resources), which was passaged once in Vero E6 cells followed by once in Vero cells upon receipt. Vero and Vero E6 cells were cultivated in DMEM containing 5% FBS and 1% penicillin-streptomycin and maintained at 37°C and 5% CO_2_.

### Animal studies

All animal studies were performed in two independent experiments. Three-to six-week-old golden Syrian hamsters (Envigo) were inoculated intranasally with 5 log_10_ PFU of SARS-CoV-2 in 100 μL PBS. Hamsters were weighed and monitored for clinical signs of illness daily. Air samples, oral swabs, nasal washes, fur swabs, and rectal swabs were collected daily. At the termination of studies, or when clinical signs of disease such as 15% weight loss were present, hamsters were euthanized via CO_2_ inhalation, and bronchoalveolar lavage (BAL) fluid and lung tissue were collected. Relative humidity and temperature in the animal facility during the studies were 27.1 ± 11% and 24.6 ± 0.8°C, respectively.

### Aerosol sampling and measurement

#### Chamber and nosecone

Aerosols generated by hamsters were sampled using two different approaches. In the first, infected hamsters were placed in a 2L sealed chamber, in which they were allowed to move freely. Air was supplied through an inlet to the chamber, and aerosols were sampled through an outlet. This approach captured total aerosols produced in exhaled breath, released from the fur, and resuspended from the chamber floor by the hamster’s activity. The second approach captured aerosols produced in exhaled breath only. Infected hamsters were anesthetized with a mixture of ketamine (100 mg/kg) and xylazine (10 mg/kg) administered by intraperitoneal (i.p.) injection. After the hamster was fully immobile, a non-rebreather nosecone (Kent Scientific, VetFlo-0802) was placed on the hamster. Air was provided through the nosecone’s inlet tube, and aerosols were sampled via the outlet port.

#### Infectious virus collection

A condensation sampler (Aerosol Devices Inc. Series 110A) was connected to the outlet of the chamber or nosecone. The sampler was operated at 1.5 L/min for a total of 1 hour, collecting aerosols directly into a vial containing 400μl BA-1 medium. To separate aerosols by size, a cyclone (URG 2000-30E-5-2.5-S) that removes aerosols larger than 8 μm at the sampling flow rate used here was installed upstream of the condensation sampler. Samples were collected for 30 minutes with the cyclone and 30 minutes without the cyclone. For 4 hamsters, the cyclone was used during the first 30-minute period, and for 4 hamsters, the cyclone was used during the second 30-minute period.

#### Aerosol size distribution

An aerodynamic particle sizer (TSI 3321) was connected to the outlet of the chamber or nosecone to measure the aerosol size distribution for 15 minutes in 1 minute intervals at a flow rate of 1 L/min, with makeup air provided at the same flow rate. This instrument detects aerosols over the size range 0.5-20 μm. Background concentrations measured in an empty chamber and nosecone were subtracted to estimate the contribution from the hamsters alone.

### Viral quantification

#### Plaque assays

Infectious virus was quantified via Vero cell plaque assay. Briefly, samples were serially diluted, plated onto confluent Vero cells in a 6-well plate, and incubated for 1 hour at 37°C. After a 1-hour adsorption, 2 mL of an 0.8% agarose overlay medium was added to the wells. Plates were incubated for 1 day at 37°C, after which a second 2 mL overlay containing 3% neutral red was added to the wells. One day later, plaques were counted. The limit of detection was 1.4 log_10_ PFU/swab (oral/fur/rectal) and BAL fluid, 0.3 log_10_ PFU/air sample, and 0.7 log_10_ PFU/nasal wash.

#### Real-time RT-PCR

Viral RNA was extracted from samples via Qiagen QIAamp Viral RNA Mini kit. RNA was quantified by real-time RT-PCR using the 2019-nCoV RUO primer/probe kit (IDT 10006713) and BioRad iTaq Universal Probes One-Step kit. Synthetic SARS-CoV-2 RNA (BEI NR-52358) was used for a standard curve. The limit of detection was 1.33 log_10_ RNA copies/oral swab, 1.66 log_10_ RNA copies/air sample, and 1.38 log_10_ RNA copies/nasal wash.

### Statistics

Samples were compared using a t-test or mixed-effects analysis with Sidak’s correction for multiple comparisons. All statistics were performed in GraphPad Prism 9.

### Ethics statement

All animal experiments were approved by the Institutional Biosafety Committee and Institutional Animal Care and Use Committee at Virginia Polytechnic Institute and State University (IACUC protocol 20-184). All experiments involving infectious SARS-CoV-2 were performed in BSL3 and ABSL3 containment.

## Results

### Aerosol generation by hamsters

To test whether inoculated animals shed SARS-CoV-2 in aerosols, we established two aerosol sampling methods for hamsters. In the first method, animals were allowed to move freely within an empty 2L chamber (Figure 1A). In the second method, animals were anesthetized, and a nosecone was placed over the nose and mouth (Figure 1B). Aerosols were sampled via an outlet port from the chamber or nosecone.

**Figure 1.**
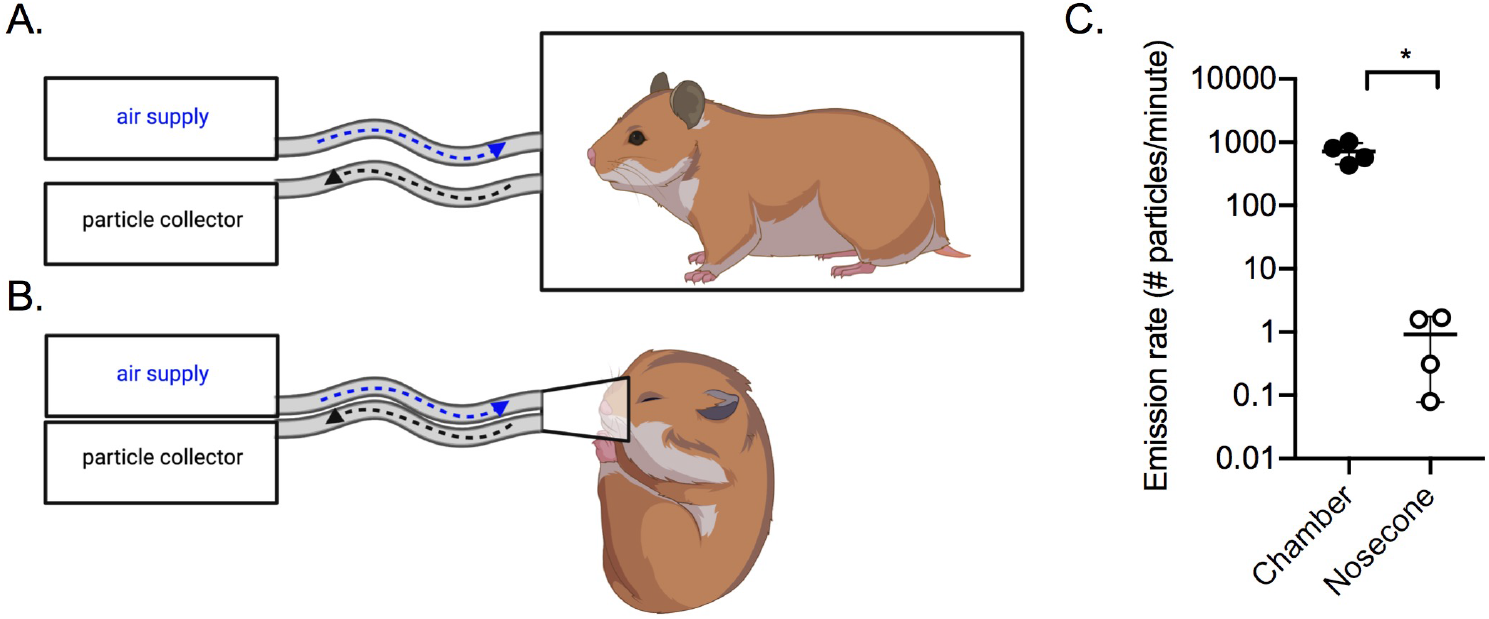
Air sampling methods. Aerosols were collected using a condensation sampler or an aerodynamic particle sizer. (A) Air was collected from awake hamsters within a sealed 2L chamber; (B) Air was collected from anesthetized hamsters using a nosecone. (C) Aerosol particle emission rate from uninfected hamsters (n=4) in the chamber (filled circles) or by nosecone (open circles). *p<0.05.

We measured aerosols produced by uninfected hamsters and calculated the emission rate. Using the chamber, we found an average of 700 aerosol particles emitted per minute per hamster (Figure 1C), with 99.9% of them <10 µm in size (Supplementary Figure 1). Using the nosecone, we found an average of 1 aerosol particle emitted per minute per hamster, which was significantly fewer than the chamber approach (p<0.05). The size distribution was similar in both cases, with very small particles being the most abundant.

### Infectious SARS-CoV-2 is emitted in aerosols

Hamsters were inoculated intranasally with SARS-CoV-2 strain USA-WA1/2020. Mild weight loss occurred from days post-inoculation (dpi) 2 through 5 (Figure 2A). Oral swabs and nasal washes were collected daily. Virus peaked on dpi 1 in the oral swabs at 3.5 log_10_ PFU/swab and on dpi 2 in the nasal washes at 3.9 log_10_ PFU/wash (Figure 2B and C). Significantly higher viral titers were observed in nasal washes from males compared to females on dpi 1, with a 5,000-fold difference (p<0.001). Viral titers for females peaked one day later than males. Viral titers decreased over time, with infectious virus below the limit of detection by dpi 5 for most animals. Samples were tested for RNA, which was detectable through dpi 10 (Figure 2E and F). Sex-specific differences were not observed for viral RNA levels.

**Figure 2.**
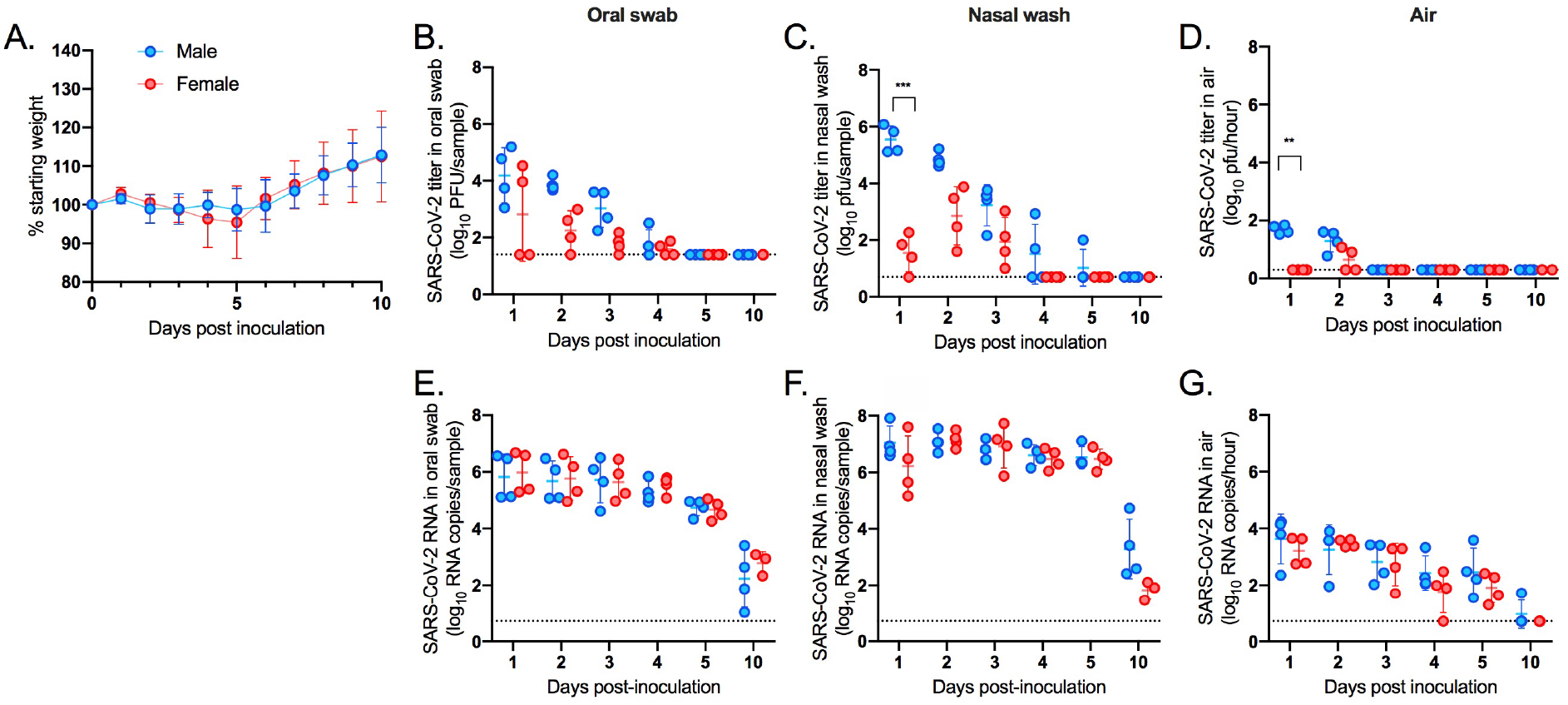
Shedding of SARS-CoV-2 in aerosols. Male (n=4) and female (n=4) hamsters were inoculated intranasally with SARS-CoV-2. (A) Percent starting weight; (B) Viral titers in oral swabs; (C) Viral titers in nasal washes; (D) Viral titers in air samples; (E) Viral RNA levels in oral swabs; (F) Viral RNA levels in nasal washes; (G) Viral RNA levels in air samples. Dashed line represents limit of detection. **p<0.01; ***p<0001.

Air samples were collected daily for 1 hour using a condensation sampler, which maintains viral infectivity [27]. Infectious virus was detected in aerosols collected on dpi 1 and 2 from males, with a mean emission rate of 1.5 log_10_ PFU/hour, and infectious virus was detected on dpi 2 from females, with a mean emission rate of 1.0 log_10_ PFU/hour (Figure 2D). Significantly greater infectious viral titers were detected in air samples from males compared to females on dpi 1 (p<0.01). The overall mean emission rate across dpi 1 and 2 was 1.4 log_10_ PFU/hour. A majority (75%) of inoculated animals released detectable levels of virus in the air on dpi 2, and emission rates ranged from 0.9 to 1.8 log_10_ PFU/hour. Infectious virus was below the limit of detection in air samples collected after dpi 2. Air samples were also tested for viral RNA; viral RNA was detected through dpi 5, with levels below the limit of detection by dpi 10 (Figure 2G). Sex-specific differences were not observed for viral RNA levels in the air. For samples with detectable infectious virus, the RNA levels were approximately 200-fold higher than PFU levels on dpi 1 and 2 for males and approximately 300-fold higher than PFU levels for females on dpi 2. Together, these data show that infectious SARS-CoV-2 is emitted in aerosols early in infection, prior to and concurrent with the onset of mild clinical signs of disease.

### Aerosols <8 µm contain infectious SARS-CoV-2

Once we established the infectious window for airborne virus, we inoculated additional hamsters in order to test whether small aerosols contain infectious SARS-CoV-2. Here, we shortened the air sampling time to 30 minutes. Male hamsters were used, as virus was more readily detected in their air samples compared to females’ air samples. To test whether the virus detected in the air was residual inoculum, we collected an air sample at 4 hours post-inoculation using the chamber; no virus was detected (Figure 3A). Then, we collected air samples on dpi 1 and 2 in the chamber, with and without a cyclone separator, which removed aerosols >8 µm, placed upstream of the condensation sampler. We detected similar titers in samples collected with and without the cyclone separator, with no statistical significance in the difference in mean titers on dpi 1 (p=0.34) or dpi 2 (p=0.37), indicating that size restriction did not alter the amount of virus detected (Figure 3A). To test whether airborne SARS-CoV-2 was detectable in the breath, we anesthetized the hamsters and collected their breath from the nosecone for one hour. Infectious virus was not detectable. However, the breath rate was very low during anesthesia (13 ± 2 breaths/minute), and very few total respiratory aerosols were collected with this method compared to the chamber method (Figure 1C).

**Figure 3.**
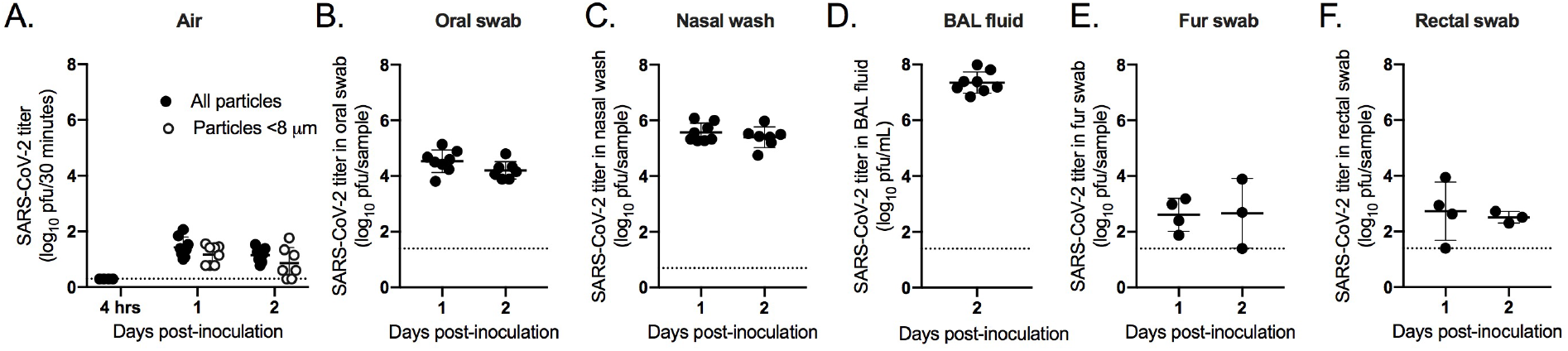
Aerosols <8 µm contain SARS-CoV-2. Male hamsters (n=8) were inoculated intranasally with SARS-CoV-2. (A) Viral titers in total aerosols (filled circles) or those <8 µm (open circles); (B) Viral titers in oral swabs; (C) Viral titers in nasal washes; (D) Viral titers in BAL fluid; (E) Viral titers in fur swabs; (F) Viral titers in rectal swabs. Dashed line represents limit of detection.

High levels of virus were detectable in oral swabs, nasal washes, and BAL fluid collected from the animals (Figure 3B-D). A low level of virus was detected in fur and rectal swabs taken from the animals, which indicates that some of the airborne virus detected in the chamber could be resuspended from the body (Figure 3E and F). Together, these results indicate that SARS-CoV-2 is emitted primarily in small aerosols <8 µm.

## Discussion

In this study, we found that hamsters emitted infectious SARS-CoV-2 in aerosols primarily <8 µm in size in the absence of severe disease, with an emission rate of 1.4 log_10_ PFU/hour (Figures 2 and 3). Peak emission of infectious virus was delayed by 1 day for females compared to males. SARS-CoV-2 viral RNA was detected in aerosols and the upper respiratory tract for a longer duration than infectious virus. The upper and lower respiratory tract contained high titers of virus, suggesting that the virus isolated from the air was primarily derived from the breath. However, we found that the fur was contaminated with low levels of infectious virus, indicating that virus resuspended from the fur may also be a viable mechanism of SARS-CoV-2 emission into the air.

Aerosols <8 µm in size can remain airborne and be inhaled. Thus, our results suggest that airborne transmission is likely a major driver of SARS-CoV-2 transmission. Our studies support reports of infectious SARS-CoV-2 collected from aerosols near COVID-19 patients [4-7], as well as studies showing non-contact transmission of SARS-CoV-2 between ferrets and hamsters [17, 18, 22], including one study that demonstrated transmission over a 1-meter distance [28]. Some previous studies have been unable to culture virus from air samples from COVID-19 patients or inoculated non-human primates due to unknown collection times post-onset of disease or the use of air sampling equipment or buffers that do not maintain viral infectivity [16, 29]. Our results support the idea that transmission is likely to occur prior to or concurrent with symptom onset or in the absence of clinical disease, supporting studies that have found that asymptomatic infection is a major driver of community transmission [23, 24]. Identifying the modes of SARS-CoV-2 transmission is critical to designing interventions to effectively prevent transmission, and our results support the use of masks and ventilation to reduce SARS-CoV-2 transmission.

This study was limited by our inability to collect EBC from the hamsters. We were unable to detect infectious virus directly from the breath when anesthetized; however, anesthesia decreased the breath rate of hamsters, and we detected 700-fold fewer aerosol particles with this method (Figure 1). Thus, given the detection of infectious virus on the fur, we cannot exclude that virus resuspended from the fur during movement may have contributed to the particles that we detected using the chamber method. Movement in guinea pigs has been shown to increase influenza particles in the air by increasing the resuspension of dust particles from the body [30]. The contribution of resuspended aerosols to SARS-CoV-2 airborne transmission has not been studied.

Sex-specific differences in COVID-19 disease severity are widely reported, with more severe disease in men [31, 32]. Here, we detected a delay in transmission potential in female hamsters compared to male hamsters. It is unclear whether these sex-specific differences in infectious viral shedding detected are relevant to human transmission. Interestingly, viral RNA levels in air samples were not significantly different between male and female hamsters, suggesting that a female-specific factor may enhance the release of defective virions. Most studies detect viral RNA, which may lead to the underestimation of differences in viral transmission potential between sexes. Sex-specific differences in infectious viral titers have not been observed by other groups, but differences in timing of samples collected may be relevant, as the largest difference was observed on dpi 1 in this study, which was not tested in other studies [33, 34].

Public health agencies have recently begun describing SARS-CoV-2 as an airborne virus. Here, we show that infectious virus is indeed culturable from the air early after infection, with the majority of aerosols containing infectious virus <8 µm in size. This suggests that SARS-CoV-2 may be maintained in the air for hours and over larger distances that previously recognized, with ventilation being an important tool for preventing transmission. Future studies will be critical for establishing the transmission potential of small aerosols containing SARS-CoV-2.

## Acknowledgements

Funding for this project was provided by a Virginia Tech Center for Emerging, Zoonotic, and Arthropod-borne Pathogens (CeZAP) seed grant and a Virginia Tech Institute for Critical Technology and Applied Science (ICTAS) Junior Faculty Award. Support was also provided by the Virginia Tech Fralin Life Sciences Institute and the BIOTRANS Interdisciplinary Graduate Education Program. We thank Dr. Amy Rizzo and TRACSS staff for contributions to the development of the animal protocol and animal husbandry. The following reagents were obtained through BEI Resources, NIAID, NIH: SARS-CoV-2 isolate USA-WA1/2020, NR-52281 and Quantitative Synthetic RNA from SARS-CoV-2, NR-52538. Figure 1 was created with BioRender.com.

---

**Supplementary Figure 1.**
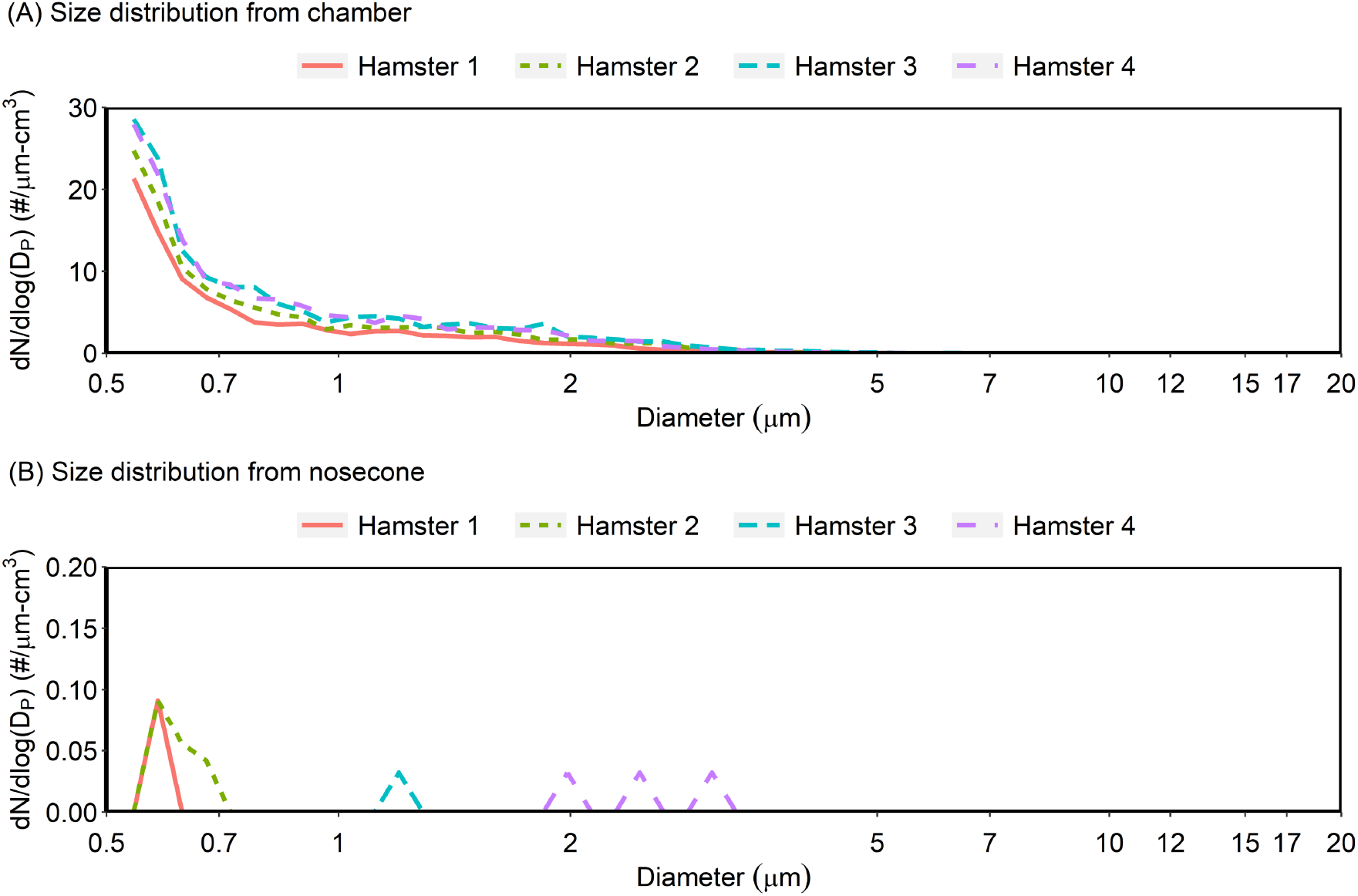
Aerosol size distributions. Size distributions of aerosols generated from uninfected hamsters (A) in the chamber or (B) through a nosecone. Each size distribution curve was obtained at times corresponding to the highest aerosol particle concentration during the 15-minute sampling period for each hamster.

## Notes

### Competing Interest Statement

The authors have declared no competing interest.

## References

1. Organization WH. WHO Coronavirus (COVID-19) Dashboard [July 2021]. Available from: https://covid19.who.int/.

2. Marr LC, Tang JW, Van Mullekom J, Lakdawala SS. Mechanistic insights into the effect of humidity on airborne influenza virus survival, transmission and incidence. J R Soc Interface. 2019;16(150):20180298. Epub 2019/04/09. doi: 10.1098/rsif.2018.0298. PubMed PMID: 30958176; PubMed Central PMCID: PMCPMC6364647.

3. L. Morawska GRJ, Z.D. Ristovski, M. Hargreaves, K. Mengersen, S. Corbett, C.Y.H. Chao, Y. Li, D. Katoshevski,. Size distribution and sites of origin of droplets expelled from the human respiratory tract during expiratory activities,. Journal of Aerosol Science. 2009;40(3):256–26. doi: 10.1016/j.jaerosci.2008.11.002.

4. Lednicky JA, Lauzardo M, Hugh Fan Z, Jutla A, Tilly TB, Gangwar M, et al. Viable SARS-CoV-2 in the air of a hospital room with COVID-19 patients. Int J Infect Dis. 2020. Epub 2020/09/20. doi: 10.1016/j.ijid.2020.09.025. PubMed PMID: 32949774; PubMed Central PMCID: PMCPMC7493737.

5. Santarpia JL, Rivera DN, Herrera VL, Morwitzer MJ, Creager HM, Santarpia GW, et al. Aerosol and surface contamination of SARS-CoV-2 observed in quarantine and isolation care. Sci Rep. 2020;10(1):12732. Epub 2020/07/31. doi: 10.1038/s41598-020-69286-3. PubMed PMID: 32728118; PubMed Central PMCID: PMCPMC7391640.

6. Guo ZD, Wang ZY, Zhang SF, Li X, Li L, Li C, et al. Aerosol and Surface Distribution of Severe Acute Respiratory Syndrome Coronavirus 2 in Hospital Wards, Wuhan, China, 2020. Emerg Infect Dis. 2020;26(7):1583–91. Epub 2020/04/11. doi: 10.3201/eid2607.200885. PubMed PMID: 32275497; PubMed Central PMCID: PMCPMC7323510.

7. Liu Y, Ning Z, Chen Y, Guo M, Liu Y, Gali NK, et al. Aerodynamic analysis of SARS-CoV-2 in two Wuhan hospitals. Nature. 2020;582(7813):557–60. Epub 2020/04/28. doi: 10.1038/s41586-020-2271-3. PubMed PMID: 32340022.

8. Lednicky JA, Lauzardo M, Alam MM, Elbadry MA, Stephenson CJ, Gibson JC, et al. Isolation of SARS-CoV-2 from the air in a car driven by a COVID patient with mild illness. Int J Infect Dis. 2021;108:212–6. Epub 2021/04/27. doi: 10.1016/j.ijid.2021.04.063. PubMed PMID: 33901650; PubMed Central PMCID: PMCPMC8064821.

9. Leung NHL, Chu DKW, Shiu EYC, Chan KH, McDevitt JJ, Hau BJP, et al. Respiratory virus shedding in exhaled breath and efficacy of face masks. Nat Med. 2020;26(5):676–80. Epub 2020/05/07. doi: 10.1038/s41591-020-0843-2. PubMed PMID: 32371934.

10. Yan J, Grantham M, Pantelic J, Bueno de Mesquita PJ, Albert B, Liu F, et al. Infectious virus in exhaled breath of symptomatic seasonal influenza cases from a college community. Proc Natl Acad Sci U S A. 2018;115(5):1081–6. Epub 2018/01/20. doi: 10.1073/pnas.1716561115. PubMed PMID: 29348203; PubMed Central PMCID: PMCPMC5798362.

11. Milton DK, Fabian MP, Cowling BJ, Grantham ML, McDevitt JJ. Influenza virus aerosols in human exhaled breath: particle size, culturability, and effect of surgical masks. PLoS Pathog. 2013;9(3):e1003205. Epub 2013/03/19. doi: 10.1371/journal.ppat.1003205. PubMed PMID: 23505369; PubMed Central PMCID: PMCPMC3591312.

12. Tovey ER, Stelzer-Braid S, Toelle BG, Oliver BG, Reddel HK, Willenborg CM, et al. Rhinoviruses significantly affect day-to-day respiratory symptoms of children with asthma. J Allergy Clin Immunol. 2015;135(3):663–9 e12. Epub 2014/12/06. doi: 10.1016/j.jaci.2014.10.020. PubMed PMID: 25476729; PubMed Central PMCID: PMCPMC7173323.

13. Zhou L, Yao M, Zhang X, Hu B, Li X, Chen H, et al. Detection of SARS-CoV-2 in Exhaled Breath from COVID-19 Patients Ready for Hospital Discharge. medRxiv. 2020:2020.05.31.20115196. doi: 10.1101/2020.05.31.20115196.

14. Ma J, Qi X, Chen H, Li X, Zhang Z, Wang H, et al. Coronavirus Disease 2019 Patients in Earlier Stages Exhaled Millions of Severe Acute Respiratory Syndrome Coronavirus 2 Per Hour. Clinical Infectious Diseases. 2020. doi: 10.1093/cid/ciaa1283.

15. Edwards DA, Ausiello D, Salzman J, Devlin T, Langer R, Beddingfield BJ, et al. Exhaled aerosol increases with COVID-19 infection, age, and obesity. Proc Natl Acad Sci U S A. 2021;118(8). Epub 2021/02/11. doi: 10.1073/pnas.2021830118. PubMed PMID: 33563754; PubMed Central PMCID: PMCPMC7923364.

16. Zhang C, Guo Z, Zhao Z, Wang T, Li L, Miao F, et al. SARS-CoV-2 Aerosol Exhaled by Experimentally Infected Cynomolgus Monkeys. Emerg Infect Dis. 2021;27(7):1979–81. Epub 2021/06/22. doi: 10.3201/eid2707.203948. PubMed PMID: 34152969.

17. Sia SF, Yan LM, Chin AWH, Fung K, Choy KT, Wong AYL, et al. Pathogenesis and transmission of SARS-CoV-2 in golden hamsters. Nature. 2020;583(7818):834–8. Epub 2020/05/15. doi: 10.1038/s41586-020-2342-5. PubMed PMID: 32408338; PubMed Central PMCID: PMCPMC7394720.

18. Chan JF, Zhang AJ, Yuan S, Poon VK, Chan CC, Lee AC, et al. Simulation of the clinical and pathological manifestations of Coronavirus Disease 2019 (COVID-19) in golden Syrian hamster model: implications for disease pathogenesis and transmissibility. Clin Infect Dis. 2020. Epub 2020/03/28. doi: 10.1093/cid/ciaa325. PubMed PMID: 32215622; PubMed Central PMCID: PMCPMC7184405.

19. Imai M, Iwatsuki-Horimoto K, Hatta M, Loeber S, Halfmann PJ, Nakajima N, et al. Syrian hamsters as a small animal model for SARS-CoV-2 infection and countermeasure development. Proc Natl Acad Sci U S A. 2020;117(28):16587–95. Epub 2020/06/24. doi: 10.1073/pnas.2009799117. PubMed PMID: 32571934; PubMed Central PMCID: PMCPMC7368255.

20. Chak-Yiu Lee A, Zhang AJ, Fuk-Woo Chan J, Li C, Fan Z, Liu F, et al. Oral SARS-CoV-2 inoculation establishes subclinical respiratory infection with virus shedding in golden Syrian hamsters. Cell Rep Med. 2020:100121. Epub 2020/09/29. doi: 10.1016/j.xcrm.2020.100121. PubMed PMID: 32984855; PubMed Central PMCID: PMCPMC7508015.

21. Rosenke K, Meade-White K, Letko MC, Clancy C, Hansens F, Liu Y, et al. Defining the Syrian hamster as a highly susceptible preclinical model for SARS-CoV-2 infection. bioRxiv. 2020. Epub 2020/10/01. doi: 10.1101/2020.09.25.314070. PubMed PMID: 32995767; PubMed Central PMCID: PMCPMC7523093.

22. Chan JF, Yuan S, Zhang AJ, Poon VK, Chan CC, Lee AC, et al. Surgical mask partition reduces the risk of non-contact transmission in a golden Syrian hamster model for Coronavirus Disease 2019 (COVID-19). Clin Infect Dis. 2020. Epub 2020/05/31. doi: 10.1093/cid/ciaa644. PubMed PMID: 32472679; PubMed Central PMCID: PMCPMC7314229.

23. Johansson MA, Quandelacy TM, Kada S, Prasad PV, Steele M, Brooks JT, et al. SARS-CoV-2 Transmission From People Without COVID-19 Symptoms. JAMA Netw Open. 2021;4(1):e2035057. Epub 2021/01/08. doi: 10.1001/jamanetworkopen.2020.35057. PubMed PMID: 33410879; PubMed Central PMCID: PMCPMC7791354.

24. Subramanian R, He Q, Pascual M. Quantifying asymptomatic infection and transmission of COVID-19 in New York City using observed cases, serology, and testing capacity. Proc Natl Acad Sci U S A. 2021;118(9). Epub 2021/02/12. doi: 10.1073/pnas.2019716118. PubMed PMID: 33571106; PubMed Central PMCID: PMCPMC7936345.

25. Iwasaki S, Fujisawa S, Nakakubo S, Kamada K, Yamashita Y, Fukumoto T, et al. Comparison of SARS-CoV-2 detection in nasopharyngeal swab and saliva. J Infect. 2020;81(2):e145–e7. Epub 2020/06/07. doi: 10.1016/j.jinf.2020.05.071. PubMed PMID: 32504740; PubMed Central PMCID: PMCPMC7270800interests.

26. Zhu J, Guo J, Xu Y, Chen X. Viral dynamics of SARS-CoV-2 in saliva from infected patients. J Infect. 2020;81(3):e48–e50. Epub 2020/07/01. doi: 10.1016/j.jinf.2020.06.059. PubMed PMID: 32593658; PubMed Central PMCID: PMCPMC7316041.

27. Lednicky JA, Lauzardo M, Fan ZH, Jutla A, Tilly TB, Gangwar M, et al. Viable SARS-CoV-2 in the air of a hospital room with COVID-19 patients. medRxiv. 2020. Epub 2020/08/15. doi: 10.1101/2020.08.03.20167395. PubMed PMID: 32793914; PubMed Central PMCID: PMCPMC7418726.

28. Kutter JS, de Meulder D, Bestebroer TM, Lexmond P, Mulders A, Richard M, et al. SARS-CoV and SARS-CoV-2 are transmitted through the air between ferrets over more than one meter distance. Nat Commun. 2021;12(1):1653. Epub 2021/03/14. doi: 10.1038/s41467-021-21918-6. PubMed PMID: 33712573; PubMed Central PMCID: PMCPMC7955093.

29. Ong SWX, Tan YK, Coleman KK, Tan BH, Leo YS, Wang DL, et al. Lack of viable severe acute respiratory coronavirus virus 2 (SARS-CoV-2) among PCR-positive air samples from hospital rooms and community isolation facilities. Infect Control Hosp Epidemiol. 2021:1–6. Epub 2021/01/26. doi: 10.1017/ice.2021.8. PubMed PMID: 33487210; PubMed Central PMCID: PMCPMC7870907.

30. Asadi S, Gaaloul Ben Hnia N, Barre RS, Wexler AS, Ristenpart WD, Bouvier NM. Influenza A virus is transmissible via aerosolized fomites. Nat Commun. 2020;11(1):4062. Epub 2020/08/20. doi: 10.1038/s41467-020-17888-w. PubMed PMID: 32811826; PubMed Central PMCID: PMCPMC7435178.

31. Peckham H, de Gruijter NM, Raine C, Radziszewska A, Ciurtin C, Wedderburn LR, et al. Male sex identified by global COVID-19 meta-analysis as a risk factor for death and ITU admission. Nat Commun. 2020;11(1):6317. Epub 2020/12/11. doi: 10.1038/s41467-020-19741-6. PubMed PMID: 33298944; PubMed Central PMCID: PMCPMC7726563.

32. GlobalHealth5050. The Sex, Gender and COVID-19 Project [July 2021]. Available from: https://globalhealth5050.org/the-sex-gender-and-covid-19-project/.

33. Dhakal S, Ruiz-Bedoya CA, Zhou R, Creisher PS, Villano JS, Littlefield K, et al. Sex Differences in Lung Imaging and SARS-CoV-2 Antibody Responses in a COVID-19 Golden Syrian Hamster Model. mBio. 2021:e0097421. Epub 2021/07/14. doi: 10.1128/mBio.00974-21. PubMed PMID: 34253053.

34. Rosenke K, Meade-White K, Letko M, Clancy C, Hansen F, Liu Y, et al. Defining the Syrian hamster as a highly susceptible preclinical model for SARS-CoV-2 infection. Emerg Microbes Infect. 2020;9(1):2673–84. Epub 2020/12/01. doi: 10.1080/22221751.2020.1858177. PubMed PMID: 33251966; PubMed Central PMCID: PMCPMC7782266.

